# Prostaglandin E_2_ directly inhibits the conversion of inducible regulatory T cells through EP2 and EP4 receptors via antagonizing TGF-β signalling

**DOI:** 10.1101/2021.04.19.440391

**Authors:** Marie Goepp, Siobhan Crittenden, You Zhou, Adriano G Rossi, Shuh Narumiya, Chengcan Yao

## Abstract

**Background and Purpose:** Regulatory T (Treg) cells are essential for control of inflammatory processes by suppressing Th1 and Th17 cells. The bioactive lipid mediator prostaglandin E_2_ (PGE_2_) promotes inflammatory Th1 and Th17 cells and exacerbates T cell-mediated autoimmune diseases. However, the actions of PGE_2_ on the development and function of Treg cells, particularly under inflammatory conditions, are debated. In this study, we examined whether PGE_2_ had a direct action on T cells to modulate *de novo* differentiation of Treg cells.

**Experimental Approach:** We employed an *in vitro* T cell culture system of TGF-β-dependent Treg induction from naïve T cells. PGE_2_ and selective agonists for its receptors, and other small molecular inhibitors were used. Mice with specific lack of EP4 receptors in T cells were used to assess Treg cell differentiation *in vivo*. Human peripheral blood T cells from healthy individuals were used to induce differentiation of inducible Treg cells.

**Key Results:** TGF-β-induced Foxp3 expression and Treg cell differentiation *in vitro* was markedly inhibited by PGE_2_, which was due to interrupting TGF-β signalling. EP2 or EP4 agonism mimicked suppression of Foxp3 expression in WT T cells, but not in T cells deficient in EP2 or EP4, respectively. Moreover, deficiency of EP4 in T cells impaired iTreg cell differentiation *in vivo*. PGE_2_ also appeared to inhibit the conversion of human iTreg cells.

**Conclusion and Implications:** Our results show a direct, negative regulation of iTreg cell differentiation by PGE_2_, highlighting the potential for selectively targeting the PGE_2_-EP2/EP4 pathway to control T cell-mediated inflammation.

**What is already known:** PGE_2_ promotes inflammatory Th1 and Th17 cells and facilitates T cell-mediated immune inflammation, but the action of PGE_2_ on Treg cells is debated.

**What does this study add:** PGE_2_ directly acts on T cells to inhibit inducible Treg cell differentiation *in vitro* and *in vivo* through its receptors EP2 and EP4 and by antagonising TGF-β signalling.

**What is the clinical significance:** Therapeutically blocking the EP4 receptor may be beneficial for management of T cell-mediated autoimmune inflammation.

## 1. Introduction

Regulatory T cells (Tregs) are a subset of T lymphocytes that play essential roles not only in the maintenance of immune homeostasis but also in the control of inflammatory responses (Sakaguchi *et al*., 2020; Savage *et al*., 2020). Treg cells actively suppress immune responses against autologous and foreign antigens *in vitro* and *in vivo*. Evidence from many mouse models and human diseases indicates that eliminating Treg cell numbers or abrogation of their functions leads to a variety of immune-mediated pathologies, including autoimmunity (e.g. multiple sclerosis, active rheumatoid arthritis, and type 1 diabetes), allergies and graft rejection (Brusko *et al*., 2005; Möttönen *et al*., 2005; M. Schneider *et al*., 2006; Viglietta *et al*., 2004; Zhang *et al*., 2008). Treg cells are characterised as expression of the surface marker CD25 (i.e. IL-2 receptor α chain, IL-2Rα) and the master transcription factor Forkhead box P3 (Foxp3) and produce the anti-inflammatory cytokine IL-10 (Sakaguchi *et al*., 2020). Foxp3 controls both Treg cell development and their unique suppressive function (Fontenot *et al*., 2003; Gavin *et al*., 2007; Hori *et al*., 2003). Loss or mutation of Foxp3 expression links to a defective development of CD4^+^CD25^+^ Treg cells, and in turn results in fatal autoimmune and inflammatory diseases, inducing a lymphoproliferative disorder in mice and leading to the IPEX (Immunodysregulation, polyendocrinopathy, enteropathy, X-linked) syndrome in human (Bennett *et al*., 2001; Brunkow *et al*., 2001).

There are two main sub-groups of Treg cells in the body: natural (nTreg) and inducible Treg (iTreg) cells. Natural Treg cells arise in the thymus and can migrate into secondary lymphoid organs (spleen, lymph nodes, etc). In addition, iTreg cells can be developed in the periphery by conversion from naïve Foxp3^−^ T effector (Teff) cells. The cytokine transforming growth factor β (TGF-β) is a regulatory cytokine with an essential role in immune responses as well as in T cell tolerance (M. O. Li *et al*., 2006; Marie *et al*., 2006). TGF-β has both a direct role in regulating T effector cell differentiation, proliferation and apoptosis and an indirect role in the maintenance of immune homeostasis (Gorelik & Flavell, 2000; Gu *et al*., 2012). It has been well documented that TGF-β is required not only for the maintenance of the suppressive function and Foxp3 expression in nTregs but also for induction of Foxp3 expression in naïve CD4^+^ T cells and convert these cells into iTregs with a regulatory phenotype (W. Chen *et al*., 2003; Fu *et al*., 2004; Shanmugasundaram R. & Selvaraj RK., 2010). Lack or blockade of TGF-β signalling reduces Treg cell numbers and impairs suppressive functions, leading to development of autoimmune diseases (Polanczyk *et al*., 2019).

Prostaglandins (PGs) are a family of bioactive lipid mediators that are generated from arachidonic acid via the activities of cyclooxygenases (COXs) and selective PG synthases (Yao & Narumiya, 2019). PGs, including PGE_2_, PGD_2_, PGF_2α_, PGI_2_, and thromboxane A_2_, play essential roles in numerous physiological and pathophysiological processes through autocrine and/or paracrine manners. Among PGs, PGE_2_ is found in the highest amounts in most tissues and is best studied. PGE_2_ has diverse effects on the development, regulation, and activity of T cells through binding to its distinct G protein–coupled receptors (called EP1-4) (Yao & Narumiya, 2019). For example, PGE_2_ inhibits T cell receptor (TCR) signalling, activation and then reduces production of cytokines such as IL-2 and IFN-*γ* through the EP2/EP4-dependent cAMP-PKA pathway (Brudvik & Taskén, 2012). However, PGE_2_ can also promote Th1 cell differentiation by inducing IL-12Rβ1 expression through EP2/EP4-dependent cAMP and PI3K signalling (Yao *et al*., 2013). Moreover, PGE_2_ also fosters IL-23-dependent Th17 cell expansion and function by inducing IL-23R expression through EP4/EP2 and the cAMP pathway (J. Lee *et al*., 2018; Yao *et al*., 2009). Importantly, emergent studies using pharmacological approaches and transgenic animal models that target PGE_2_ receptors have demonstrated that the actions of PGE_2_ on T cells promotes immune-associated chronic inflammatory diseases in rodents and humans (including multiple sclerosis, rheumatoid arthritis, inflammatory skin and gut inflammation) (Q. Chen *et al*., 2010; Esaki *et al*., 2010; J. Lee *et al*., 2018; Robb et al., 2017; Schiffmann *et al*., 2014; Yao *et al*., 2009, 2013). While PGE_2_ was initially described to facilitate iTreg cell differentiation *in vitro* (Q. Chen *et al*., 2010; Esaki *et al*., 2010; J. Lee *et al*., 2018; Robb *et al*., 2017; Schiffmann *et al*., 2014; Yao *et al*., 2009, 2013), it has also been reported to inhibit Foxp3 induction and reduce Treg cell numbers (L. Chen *et al*., 2017; H. Li *et al*., 2017; Sahin & Sahin, 2020). We have recently reported a T cell-independent function of PGE_2_ on facilitation of Foxp3^+^ Treg cell responses in the intestine (Crittenden *et al*., 2021). However, whether and how PGE_2_ directly influences iTreg cell differentiation remains to be elucidated.

In this study, we have examined the direct actions of PGE_2_ in iTreg differentiation *in vitro* and *in vivo* using mice deficient in EP2 and EP4 receptors and highly selective small molecular reagents that target the respective PGE_2_ receptors. We found that PGE_2_ negatively regulated iTreg cell differentiation *in vitro* by inhibiting TGF-β-driven Foxp3 induction through EP2 and EP4. Lack of EP4 specifically in T cells increased Treg cell generation *in vivo*. The PGE_2_ pathway also appears to inhibit human iTreg cell differentiation. Our results have revealed that PGE_2_ directly acts on T cells to abrogate iTreg cell differentiation, which may contribute to foster T cell-mediated inflammation.

## 2. Methods

### 2.1 Animals

EP2^+/+^, EP2^−/–^ (Hizaki *et al*., 1999), EP4^+/+^, EP4^−/–^ (Segi *et al*., 1998), Lck^Cre^EP4^fl/fl^ (A. Schneider *et al*., 2004; Yao *et al*., 2013), *Rag1*^−/–^, Foxp3^YFP-Cre^ (Rubtsov *et al*., 2008) and wild-type C57BL/6 mice were bred and maintained under specific pathogen-free conditions in accredited animal facilities at the University of Edinburgh and Kyoto University. Wild-type mice were purchased from Harlan UK. Age-(>7-weeks old) and sex-matched mice were used. Mice were randomly allocated into different groups and analysed individually. No mice were excluded from the analysis. All experiments were conducted in accordance with the UK Animals (Scientific Procedures) Act of 1986 with local ethical approval from the University of Edinburgh Animal Welfare and Ethical Review Body (AWERB) or approved by the Committee on Animal Research of Kyoto University Faculty of Medicine.

### 2.2 Reagents and Antibodies

16,16-dimethyl prostaglandin E_2_ (dm-PGE_2_) and PGE_2_ were purchased from Cayman Chemical. Highly selective agonists for EP1 (ONO-DI-004), EP2 (ONO-AE1-259-01), EP3 (ONO-AE-248) or EP4 (ONO-AE1-329) were gifts from Ono Pharmaceutical Co., Japan. Selective antagonists against EP2 (PF-04418948) and EP4 (L-161,982) were purchased from Cayman. Recombinant human TGF-β1 and mouse or human IL-2 were purchased from R&D system or Biolegend. Indomethacin, dibutyryl-cAMP (db-cAMP), 3-isobutyl-1-methylxanthine (IBMX), H-89, LY-294002, AS1842856, and STAT5 inhibitor were purchased from Sigma or Calbiochem.

### 2.3 T cell transfer

Naive CD4^+^CD25^−^ CD62L^hi^ T cells were prepared from spleens of EP4^fl/fl^ or Lck^Cre^EP4^fl/fl^ mice by flow cytometry cell sorting. Cells (5 × 10^5^ cells per mouse) were transferred intravenously into Rag1^−/–^ mice. Mice were culled at 6 weeks after T cell transfer. Colons were collected for *ex vivo* analysis of lamina propria leukocytes.

### 2.4 DSS application

Wild type C57BL/6 mice were given drinking water with dextran sulfate sodium (DSS, 2% w/v) or DSS plus indomethacin (5 mg per kg body weight per day) for 5 consecutive days before colons were collected for *in vitro* analysis of T cells.

### 2.5 DNFB application

EP4^fl/fl^ and Lck^Cre^EP4^fl/fl^ mice were sensitised with 25 µl of 1% (w/v) Dinitrofluorobenzene (DNFB) in acetone/olive oil (4/1, v/v) on shaved abdominal skin on day 0. Skin draining lymph node cells were collected on day 5 for *ex vivo* analysis of T cells.

### 2.6 T cell isolation and *in vitro* culture

Mouse CD4^+^CD25^-^ naïve T cells were isolated from spleens using Miltenyi Treg cell isolation kits. CD4^+^CD25^-^Foxp3(YFP)^-^ naïve T cells and CD4^+^CD25^+^Foxp3(YFP)^+^ nTreg cells were isolated from Foxp3^YFP-Cre^ mouse spleens by flow cytometry. Cells were cultured in complete RPMI1640 medium containing 10% FBS and stimulated with plate-bound anti-CD3 (5 µg/ml) and anti-CD28 (5 µg/ml) antibodies plus various cytokines (IL-2, rhTGF-β1) and other compounds as indicated for 3 days. Human CD4^+^CD45RA^-^ naïve T cells were isolated from peripheral blood of healthy individuals, stimulated with plate-bound anti-CD3 and anti-CD28, and then cultured with IL-2 (10 ng/ml) and/or rhTGF-β1 (10 ng/ml or indicated concentrations) for 3 days. PGE_2_ (1 μM or indicated concentrations) and its receptor agonists (1 μM) and other small molecular chemicals were added at the beginning of the culture or 24 hours later. Work with human blood cells was approved by the Centre for Inflammation Research (CIR) Blood Resource (AMREC Reference number 20-HV-069).

### 2.7 Isolation of intestinal lamina propria leukocytes

Intestinal lamina propria cells were isolated as described previously (Duffin *et al*., 2016).

### 2.8 Staining and flow cytometry

For surface staining, cells were first stained with the Fixable Viability Dye eFluor® 780 (eBioscience) on ice for 30 min. After wash, cells were stained on ice for another 30 min with anti-CD45 (clone 30-F11), anti-CD3e (Clone 145-2C11), anti-CD4 (Clone GK1.5), anti-CD25 (clone PC61.5). For staining of transcription factors, cells were fixed in the Foxp3/Transcription Factor Fix Buffer (eBioscience) for 2 h or overnight followed by staining with anti-mouse Foxp3 (clone FJK-16s), anti-mouse Ki-67 (clone 16A8) for at least 1 h. For cytokine staining, cells were restimulated ex vivo with PMA (50 ng/ml) and ionomycin (750 ng/ml) for 4 h in the presence of GolgiPlug (BD Bioscience), and then fixed and permeabilised following intracellular staining with anti-mouse IL-17A (clone ReBio17B7) and anti-mouse IFN-*γ* (clone RA3-6B2) for 30 min. All Abs were purchased from eBioscience, Biolegends, or BD Bioscience. Flow cytometry was performed on the BD LSR Fortessa (BD Bioscience) and analysed by FlowJo software (Tree Star).

### 2.9 Real-time PCR

RNA purification from sorted MNPs was performed by using the RNeasy Mini Kit (Qiagen). cDNA was obtained by reverse transcription using the High-capacity cDNA Reverse Transcription Kits (ABI). Samples were analyzed by real-time PCR with LightCycler Taqman Master (Roche) and Universal ProbeLibrary (UPL) Set, Mouse (Roche) on the LightCycler 2.0 (Roche). Primers were used are Glyceraldehyde-3-phosphate dehydrogenase (*Gapdh*) forward, 5’-tgaacgggaagctcactgg-3’; *Gapdh* reverse, 5’-tccaccaccctgttgctgta-3’. *Foxp3* forward: 5′-cacccaggaaagacagcaacc-3’; *Foxp3* reverse: 5′-gcaagagctcttgtccattga-3’. *Tgfbr1* forward: 5’-aatgttacgccatgaaaatatcc-3’; *Tgfbr1* reverse: 5’-cgtccatgtcccattgtctt-3’; UPL Probe: #84. *Tgfbr2* forward: 5’-ggctctggtactctgggaaa-3’; *Tgfbr2* reverse: 5’-aatgggggctcgtaatcct-3’; UPL Probe: #7. *Smad6* forward: 5’-gttgcaacccctaccacttc-3’; *Smad6* reverse: 5’-ggaggagacagccgagaata-3’; UPL Probe: #70. *Smad7* forward: 5’-acccccatcaccttagtcg-3’; *Smad7* reverse: 5’-gaaaatccattgggtatctgga-3’; UPL Probe: #63. Expression was normalized to *Gapdh* and presented as relative expression to the control group by the 2^−ΔΔCt^ method.

### 2.10 Human gene expression analysis

We retrieved microarray data from Gene Expression Omnibus under an accession code (GSE71571) (Thomas *et al*., 2015). Raw data were normalized using the GC-RMA method (Wu *et al*., 2004). When multiple probe sets were present for a gene, the one with the largest variance was selected (Talloen *et al*., 2010). Change of the normalized expression levels for each gene by aspirin (i.e. aspirin–placebo) in colon biopsies was transformed into Z-score, which was used to estimate the alteration of PGE_2_ pathway in each patient in response to Aspirin administration. The signature score of PGE_2_ pathway was estimated using a method described previously (Bueno *et al*., 2016). Briefly, we curated a gene list representative of PGE_2_ signature including its synthases and receptors. The final list consisted of *PTGS1, PTGS2, PTGES, PTGES2, PTGES3, PTGER2* and *PTGER4*. We weighted gene expression and computed a signature score per sample using singular-value decomposition. Pearson’s correlation coefficient was used to measure the association between PGE_2_ signature and expression of Treg genes on a Z-score scale.

### 2.11 Statistical analysis

Data were expressed as mean ± SEM, and statistical significance was performed by unpaired Student’s *t* test or analysis of variance (ANOVA) with post hoc Holm-Sidak’s multiple comparisons test using Prism software (GraphPad). All *P* values <0.05 were considered as significant. Correlation analysis was calculated by Pearson’s correlation coefficient (r).

## 3. Results

### 3.1 PGE_2_ suppresses mouse iTreg differentiation *in vitro*

We firstly examined whether PGE_2_ had an impact on iTreg differentiation *in vitro*. We isolated splenic CD4^+^CD25^-^ naïve T cells from wild-type (WT) C57BL/6 mice, stimulated with anti-CD3 and anti-CD28 antibodies (Abs) and cultured with TGF-β to induce the differentiation of iTreg cells. We added different concentrations of PGE_2_ (0 to 1000 nM) at the beginning of TCR stimulation on day 0. TGF-β-induced Foxp3 expression in CD4^+^ T cells was suppressed by addition of PGE_2_ in a concentration-dependent manner (**Figure 1A, B**). To avoid PGE_2_ inhibition of TCR activation when it was added at the same time of anti-CD3 stimulation (Yao *et al*., 2013), we tested the effect of PGE_2_ by postponing its time of addition to 24 h (day 1) after anti-CD3 stimulation. Under this condition, PGE_2_ still inhibited TGF-β-induced Foxp3 expression (**Figure 1A, B**), suggesting that PGE_2_ prevents TGF-β-induced iTreg cell differentiation independently of its suppression on TCR activation.

**Figure 1.**
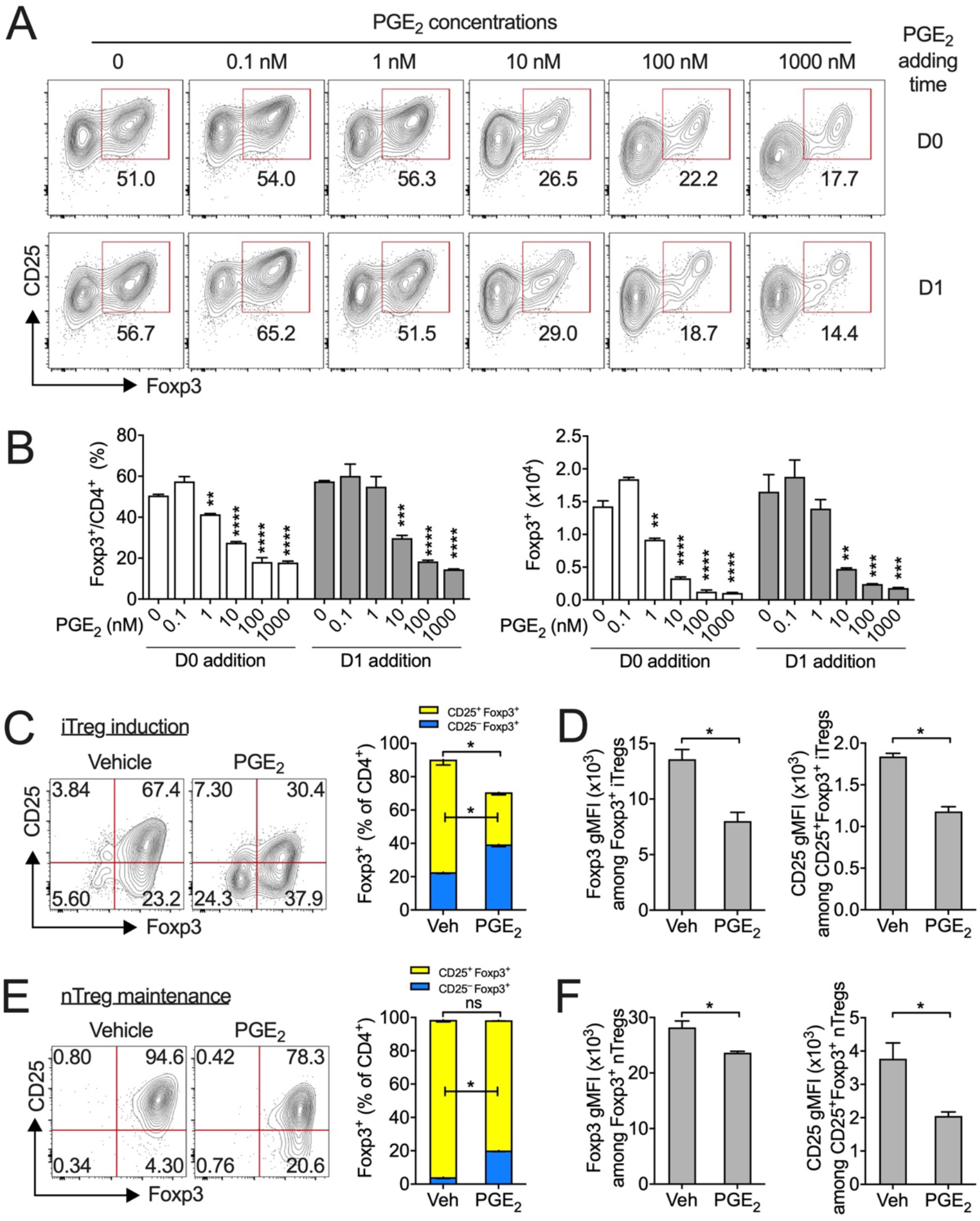
PGE_2_ suppresses iTreg cell differentiation *in vitro*. (**A**) Representative flow cytometry dot-plot of CD25 and Foxp3 expression in CD4^+^CD25^-^ naïve T cells cultured under iTreg cell priming conditions (i.e., with IL-2 and TGF-β1) for 3 days. PGE_2_ was added with indicated concentrations and at indicated time points (i.e. day 0 or 1). (**B**) Accumulated percentages and numbers of Foxp3^+^ T cells. (**C**) Representative flow cytometry dot-plot of CD25 and Foxp3 expression (left) and percentages of Foxp3^+^ T cells (right) in CD4^+^CD25^-^ Foxp3(YFP)^-^ naïve T cells cultured with IL-2 and TGF-β1 in the absence or presence of PGE_2_ for 3 days. (**D**) Geometric mean fluorescent intensity (gMFI) of Foxp3 and CD25 among Foxp3^+^ T cells. (**E)** Representative flow cytometry dot-plot of CD25 and Foxp3 expression (left) and percentages of Foxp3^+^ T cells (right) in CD4^+^CD25^-^Foxp3(YFP)^+^ nTreg cells cultured with IL-2 and TGF-β1 in the absence or presence of PGE_2_ for 3 days. (**F**) gMFI of Foxp3 and CD25 among Foxp3^+^ T cells. All experiments were performed in triplicates and repeated at least twice independently. \**P*<0.05; \*\**P*<0.01; \*\*\**P*<0.001; \*\*\*\**P*<0.0001. ns, not significant.

A very small sub-population of splenic CD4^+^CD25^-^ naïve T cells may express Foxp3. To examine whether the contamination of this small population of Foxp3^+^CD4^+^CD25^-^ T cells affects PGE_2_ inhibition on iTreg induction, we sorted splenic CD4^+^CD25^-^Foxp3(YFP)^-^ naïve T cells from Foxp3^YFP-Cre^ mice (Rubtsov *et al*., 2008) and cultured them with TGF-β. With this culture system, PGE_2_ still inhibited Foxp3 induction (**Figure 1C**). Interestingly, PGE_2_ specifically reduced CD25^+^Foxp3^+^ cells (**Figure 1C**), a Treg subpopulation that has greater immunosuppressive function compared to CD25^-^Foxp3^+^ Treg cells (Bonelli *et al*., 2009; Polanczyk *et al*., 2019). Furthermore, PGE_2_ treatment reduces mean fluorescent intensity (MFI) of Foxp3 and CD25 (**Figure 1D**), suggesting that PGE_2_ also inhibits Foxp3 expression at the single cell level.

To examine whether PGE_2_ affects the stability of Foxp3 expression on nTreg cells, we sorted splenic CD4^+^CD25^+^Foxp3(YFP)^+^ nTreg cells from Foxp3^YFP-Cre^ mice and cultured with TGF-β for 3 days. Addition of PGE_2_ did not affect total percentage of Foxp3^+^ cells, but appeared to reduce the mean fluorescence intensity (MFI) of Foxp3 (**Figure 1E, F**). Moreover, PGE_2_ treatment significantly reduced CD25 expression, leading to a reduction of the CD25^+^Foxp3^+^ nTreg subpopulation (**Figure 1E, F**). Taken together, these results suggest that PGE_2_ represses both *de novo* iTreg cell differentiation and, to a less extent, Treg maintenance.

### 3.2 EP2 and EP4 receptors mediate PGE_2_ suppression of iTreg differentiation *in vitro*

Next, we investigated which PGE_2_ receptors mediated the suppression of iTreg differentiation. We isolated splenic CD4^+^ T cells from EP2- or EP4-deficient and WT control mice, cultured with TGF-β with the addition of dm-PGE_2_ (a stable PGE_2_ analogue) or selective agonists for PGE_2_ receptors EP1-EP4. In EP2^+/+^ (on the C57BL/6 genetic background) or EP4^+/+^ mice (on the mixed C57BL/6 x 129 genetic background), EP2 and EP4 agonists mimicked PGE_2_ suppression of TGF-β-induced Foxp3 expression from CD4^+^CD25^-^ naïve T cells (**Figure 2A, C**). In EP2^-/-^ CD4^+^CD25^-^ naïve T cells, however, EP2 agonist failed to suppress Foxp3 expression although PGE_2_ and EP4 agonist still have inhibitory effects (**Figure 2B**). Likewise, EP4 agonist had no effect on induction of Foxp3 expression from EP4^-/-^ CD4^+^CD25^-^ naïve T cells, but PGE_2_ and EP2 agonist still repressed iTreg induction (**Figure 2D**). Selective EP1 and EP3 agonists appeared only mild inhibition of Foxp3 induction in C57BL/6 EP2^+/+^ T cells and had no influences on EP2^-/-^, EP4^+/+^ or EP4^-/-^ T cells (**Figure 2A-D**). Furthermore, inhibition of Foxp3 expression by PGE_2_ was rescued by combination of EP2 and EP4 antagonists, but not by blockade of either single receptor (**Figure 2E**). These results suggest that PGE_2_ suppression of iTreg cell differentiation *in vitro* is redundantly mediated by EP2 and EP4 receptors.

**Figure 2.**
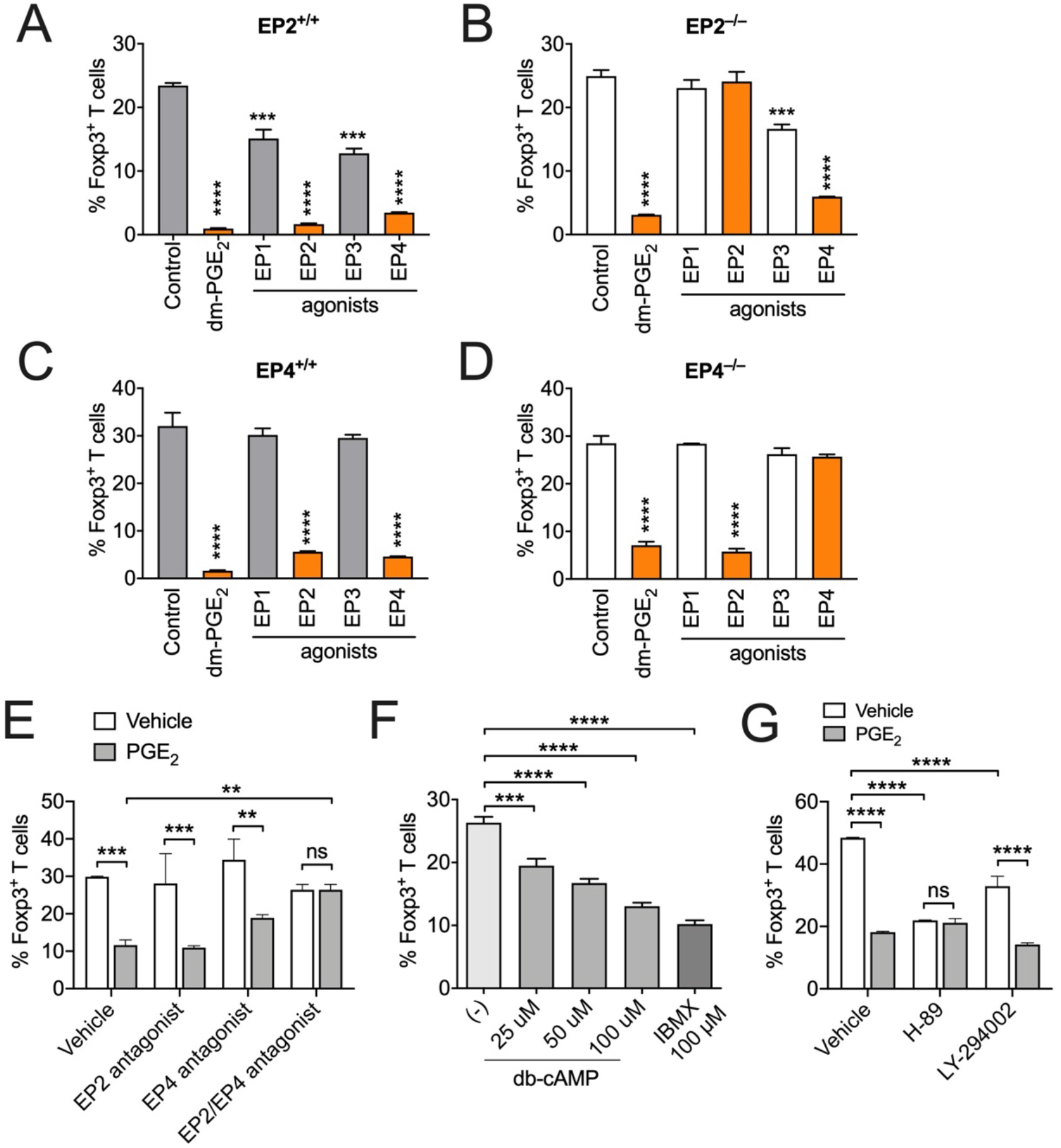
EP2 and EP4 receptors mediate PGE_2_ suppression of iTreg cell differentiation *in vitro*. (**A**,**B**) Percentages of Foxp3^+^ T cells in EP2^+/+^ (A) or EP2^-/-^ (B) CD4^+^CD25^-^ naïve T cells cultured with IL-2 and TGF-β1 with dm-PGE_2_ or selective agonists for each EP1-4 receptor for 3 days. (**C**,**D**) Percentages of Foxp3^+^ T cells in EP4^+/+^ (C) or EP4^-/-^ (D) CD4^+^CD25^-^ naïve T cells cultured with IL-2 and TGF-β1 with dm-PGE_2_ or selective agonists for each EP1-4 receptor for 3 days. (**E**) Percentages of Foxp3^+^ T cells in wild type C57BL/6 CD4^+^CD25^-^ naïve T cells cultured with IL-2 and TGF-β1 in the absence or presence of PGE_2_, EP2 antagonist or EP4 antagonist or both EP2 and EP4 antagonists for 3 days. (**F**) Percentages of Foxp3^+^ T cells in wild type C57BL/6 CD4^+^CD25^-^ naïve T cells cultured with IL-2 and TGF-β1 with db-cAMP or IBMX for 3 days. (**G**) Percentages of Foxp3^+^ T cells in wild type C57BL/6 CD4^+^CD25^-^ naïve T cells cultured with IL-2 and TGF-β1 with PGE_2_, a PKA inhibitor (H-89) or a PI3K inhibitor (LY-294002) for 3 days. All experiments were performed in triplicates and repeated at least twice independently. \**P*<0.05; \*\**P*<0.01; \*\*\**P*<0.001; \*\*\*\**P*<0.0001. ns, not significant.

Given EP2 and EP4 activate the cyclic adenosine monophosphate (cAMP) and PI3K signalling pathways (Yao & Narumiya, 2019), we examined whether these pathways mediate the suppression of iTreg cell induction. We used dibutyryl cAMP (db-cAMP, a cell-permeable cAMP analogue) and isobutylmethylxanthine (IBMX, a phosphodiesterase inhibitor that blocks cAMP degradation) to increase the intracellular cAMP levels. Similar to PGE_2_, both db-cAMP and IBMX prevented TGF-β-dependent conversion of Foxp3^+^ iTreg cell (**Figure 2F**). Blockade of the cAMP pathway by a PKA inhibitor (H-89) or the PI3K pathway by LY-294002 repressed TGF-β-dependent Foxp3 expression (**Figure 2G**). PGE_2_ had no additive suppression of Foxp3 induction with H-89, but did further reduced Foxp3 expression in the presence of LY-294002 (**Figure 2G**). These results indicate that the cAMP/PKA, rather than PI3K, pathway is involved in PGE_2_-dependent inhibition of iTreg cell differentiation.

### 3.3 PGE_2_ antagonises TGF-β signalling during iTreg differentiation

We next examined the mechanisms by which PGE_2_ inhibits iTreg cell differentiation. We stained T cells with Ki-67, an intracellular marker of cell proliferation. In the absence of TGF-β, PGE_2_ moderately prevented anti-CD3/CD28-induced naïve T cell proliferation, evidenced as Ki-67^+^Foxp3^-^ T cells (**Figure 3A**). Under the iTreg cell differentiation condition, TGF-β markedly induced and expand Ki-67^+^Foxp3^+^ proliferative iTreg cells (**Figure 3A**). However, this was significantly prevented by PGE_2_ which had few effects on Ki-67^+^Foxp3^-^ non-Treg cells (**Figure 3A**), indicating that PGE_2_ selectively prevents TGF-β-dependent induction of proliferating iTreg cells. Indeed, PGE_2_ suppressed TGF-β responsiveness during Foxp3^+^ iTreg differentiation (**Figure 3B**).

**Figure 3.**
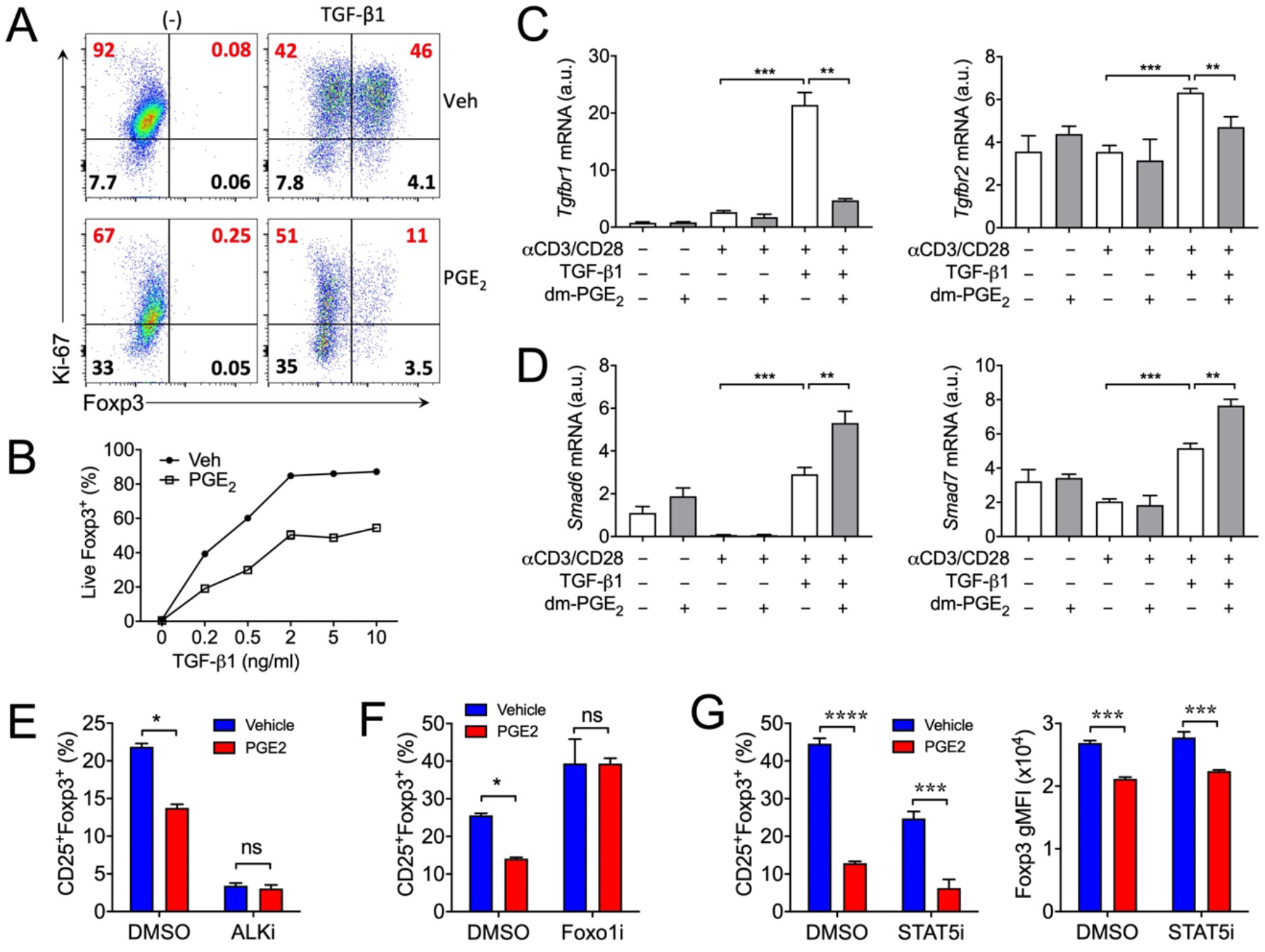
PGE_2_ antagonises TGF-β signalling during iTreg cell differentiation. (**A**) Representative flow cytometry dot-plot of Foxp3 and Ki-67 expression in CD4^+^CD25^-^ naïve T cells cultured with IL-2 and TGF-β1 in the absence or presence of PGE_2_ for 3 days. (**B**) Percentages of live Foxp3^+^ T cells in CD4^+^CD25^-^ naïve T cells cultured with IL-2 and indicated concentrations of TGF-β1 in the absence or presence of PGE_2_ for 3 days. (**C**,**D**) Expression of *Tgfbr1, Tgfbr2, Smad6* and *Smad7* genes in CD4^+^CD25^-^ naïve T cells cultured with or without anti-CD3/CD28, TGF-β1 or PGE_2_ for 3 days. (**E-G**) Percentages of CD25^+^Foxp3^+^ T cells in CD4^+^CD25^-^ naïve T cells cultured with IL-2 and TGF-β1 in the absence or presence of PGE_2_ and inhibitors for ALK (ALKi, E), Foxo1 (Foxo1i, F) or STAT5 (STAT5i, G) for 3 days. Geometric mean fluorescent intensity (gMFI) of Foxp3 among Foxp3^+^ T cells (G). \**P*<0.05; \*\**P*<0.01; \*\*\**P*<0.001. ns, not significant.

During iTreg differentiation, TGF-β firstly activates gene expression of its receptors (i.e. *Tgfbr1* and *Tgfbr2*) on T cells, which were both repressed by the addition of PGE_2_ (**Figure 3C**). TGF-β also stimulates gene expression of Smad6 and Smad7, endogenous inhibitors for TGF-β signalling, which were significantly further upregulated by PGE_2_ (**Figure 3D**). These results suggest an inhibitory effect of PGE_2_ on TGF-β signalling in T cells, as seen in other cell types (Lenicov *et al*., 2018; P. E. Thomas *et al*., 2007; Wettlaufer *et al*., 2017). To further study the possibility of PGE_2_ influence on TGF-β signalling, we used a small molecular ALK inhibitor which blocks the TGF-β/TGF-β receptor/Smad pathway. ALK inhibitor itself significantly repressed TGF-β-dependent iTreg cell induction, and addition of PGE_2_ had no additional effects on Foxp3 induction in the present of with the ALK inhibitor (**Figure 3E**). The transcription factor Foxo1 acts downstream of TGF-β receptors, and is responsible for TGF-β responsiveness in iTreg cell differentiation (Kerdiles *et al*., 2010). The Foxo1 inhibitor (AS1842856) did not affect TGF-β-dependent Foxp3 induction, but it reversed PGE_2_ suppression of Foxp3 induction (**Figure 3F**). These results suggest that PGE_2_ suppresses the process of iTreg differentiation by antagonizing TGF-β signalling.

In response to TCR engagement, activated T cells produce large amount of IL-2, which is also essential for iTreg cell differentiation through the transcription factor STAT5 (Davidson *et al*., 2007; Guo *et al*., 2013). As PGE_2_ strongly inhibits TCR activation and IL-2 production, we asked whether PGE_2_ suppresses iTreg cell induction via inhibiting IL-2-STAT5 signalling. We cultured T cells under the iTreg-skewing condition and used a STAT5 inhibitor (STAT5i). As expected, the STAT5 inhibitor suppressed iTreg cell conversion compared to vehicle control (**Figure 3G**). However, PGE_2_ was still able to further down-regulate Foxp3 expression in the presence of STAT5 inhibitor (**Figure 3G**). Thus, IL-2-STAT5 signalling is unlikely to be involved in PGE_2_ suppression of iTreg cell induction.

### 3.4 Lack of EP4 impairs iTreg cell differentiation *in vivo*

We have recently found that blockade of endogenous PGE_2_ production in naïve WT mice by inhibition of COX activities increased Foxp3^+^ Treg cell numbers in various organs (**Crittenden *et al***., **2021**). To examine whether blockade of endogenous PGE_2_ production also enhances Treg cell responses under inflammatory conditions, we used 2% DSS to induce acute colonic inflammation WT C57BL/6 mice. DSS treatment increased accumulation of total T cells in the colon, which was further enhanced by co-administration of indomethacin, a non-selective COX inhibitor (**Figure 4A**). This is consistent with previous report that blocking COX activity exacerbated DSS-dependent intestinal inflammation (**Duffin *et al*. 2016**). Interestingly, indomethacin also significantly increased numbers of Foxp3^+^ Treg cells, but not Foxp3^−^ Teff cells, in inflamed colons (**Figure 4B, C**), which was in line with upregulated Foxp3 gene expression in the colon tissues (**Figure 4D**). These results suggest that endogenous PG signalling represses Treg cell response under inflammatory conditions.

**Figure 4.**
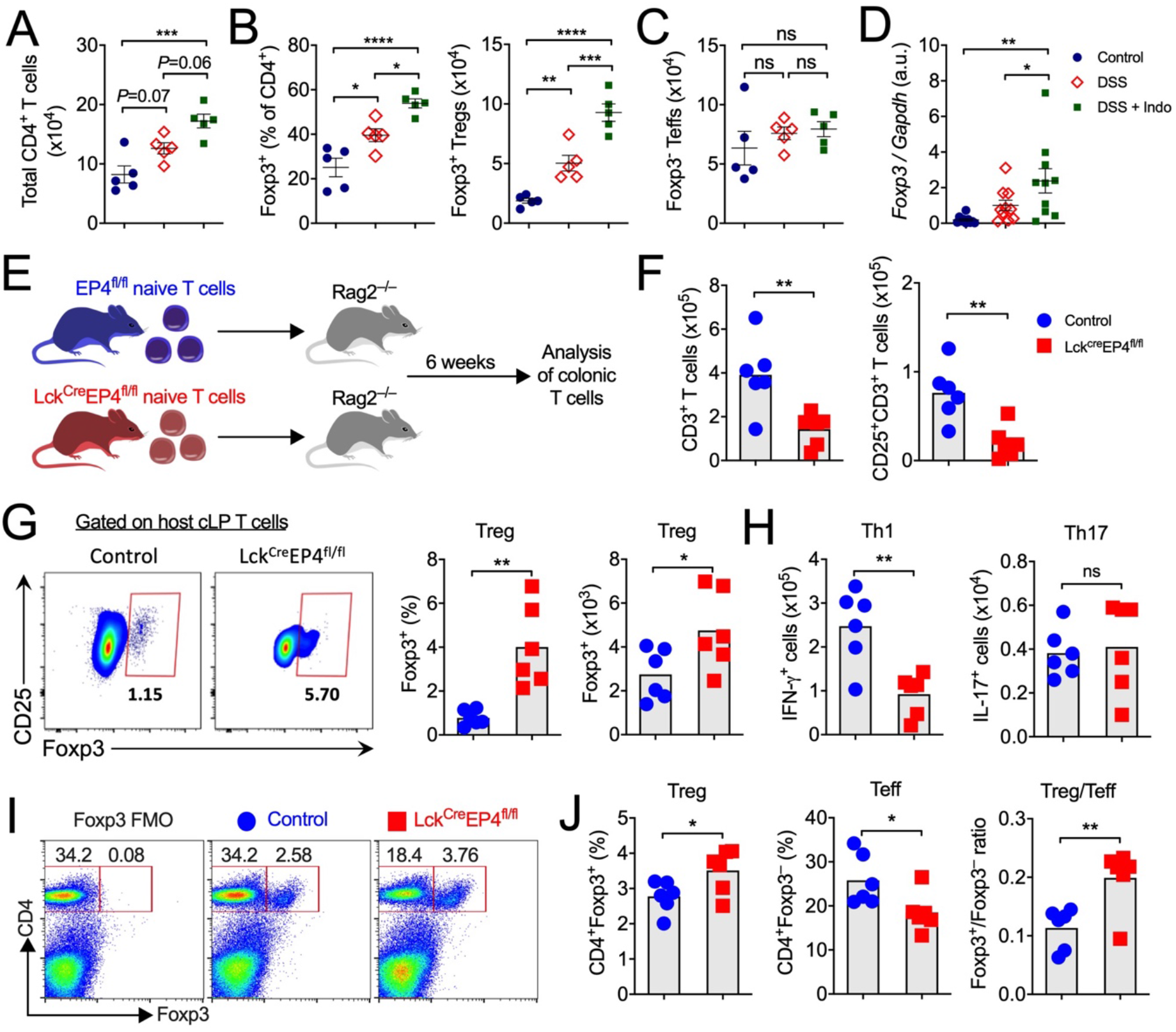
PGE_2_ represses Treg cell differentiation *in vivo*. (**A**) Total CD3^+^CD4^+^ T cells in colonic lamina propria of mice treated with vehicle or 2% DSS in drinking water or DSS plus indomethacin in drinking water for 5 days. **(B)** Percentages and numbers of colonic Foxp3+ Treg cells. **(C)** Numbers of colonic Foxp3– Teff cells. **(D)** Foxp3 gene expression in whole colon tissues. **(E)** Schematic representation of the experimental protocol for T cells transfer. CD4^+^CD25^-^CD62L^hi^ naïve T cells isolated from LcK^Cre^EP4^fl/fl^ and control EP4^fl/fl^ mice were transferred into Rag1^-/-^ mice. Colonic lamina propria T cells in host Rag1^-/-^ mice were analysed 6 weeks later. (**F**) Numbers of colonic CD3^+^ total T cells and CD25^+^ activated T cells. (**G**) Representative flow cytometry dot-plot of Foxp3 and CD25 expression, percentages and absolute numbers of Foxp3^+^ T cells in colons. (**H**) Absolute numbers of colonic Th1 and Th17 cells. (**I**) Representative flow cytometry dot-plot of Foxp3 and CD4 expression in skin draining lymph nodes of LcK^Cre^EP4^fl/fl^ and control EP4^fl/fl^ mice that were sensitised with DNFB. (**J**) Percentages of Foxp3^+^ Treg and Foxp3^-^ T effector (Teff) cells and the ratio of Treg vs Teff cells in dLNs. Each dot represents one mouse. \**P*<0.05; \*\**P*<0.01; \*\*\**P*<0.001; \*\*\*\**P*<0.0001. ns, not significant.

To further examine whether PGE_2_ signalling directly modulates Treg cell responses *in vivo*. We crossed EP4-floxed mice to Lck-Cre mice to generate T cell-specific EP4 deficient mice (Lck^Cre^EP4^fl/fl^). Lck^Cre^EP4^fl/fl^ and control EP4^fl/fl^ mice had comparable nTregs in the thymus (Crittenden *et al*., 2021), suggesting that lack of EP4 signalling in T cells does not affect nTreg cell development *in vivo*. To examine whether PGE_2_ affects iTreg cell differentiation *in vivo*, we sorted naïve CD3^+^CD4^+^CD25^-^CD62L^+^ T cells from Lck^Cre^EP4^fl/fl^ and control EP4^fl/fl^ mice, and then transferred these cells into Rag1^-/-^ mice that have no T and B cells (**Figure 4E**). Upon transfer, naïve T cells are activated, proliferated and differentiated into T effect cells (e.g. Th1 and Th17 cells) in the host mice and accumulated in the large intestines. Simultaneously, a small population of T cells are differentiated into Foxp3^+^ iTreg cells. Lack of EP4 signalling reduced total T cells migration to the colon and down-regulation of T cell activation evidenced by reduction of CD25 expression (**Figure 4F**). In contrast, differentiation of Foxp3^+^ Tregs in the host mouse colons from EP4-deficient naive T cells was greater than that from control EP4-sufficient naïve T cells (**Figure 4G**). In agreement with our previous findings (Yao *et al*., 2013), Rag1^-/-^ mice transferred with EP4-deficient naïve T cells had less IFN-*γ*^+^ Th1 cells compared to mice that were transferred with control naïve T cells, but EP4 deficiency had no influence on colonic IL-17^+^ Th17 cells in the host mice (**Figure 4H**). To further confirm the effect of EP4 signalling on Treg responses *in vivo*, we sensitised Lck^Cre^EP4^fl/fl^ and control EP4^fl/fl^ mice with a hapten dinitrobenzfluorene (DNBF) on the abdominal skin and analysed T cells in skin-draining lymph nodes. Again, lack of EP4 signalling in T cells significantly increased Foxp3^+^ Treg cells but reduced Foxp3^-^ effector T cells in draining lymph nodes (**Figure 4I, J**). Together, these results indicate that PGE_2_-EP4 signalling directly acts on T cells to impede iTreg cell differentiation *in vivo*.

### 3.5 Inhibition of human iTreg cell differentiation by PGE_2_

To corroborate our findings from mouse T cells, we examined whether PGE_2_ suppresses human iTreg cell differentiation. We isolated CD4^+^CD45RA^-^ naïve T cells from peripheral blood of healthy individuals, stimulated with anti-CD3 and anti-CD28 Abs, and cultured with IL-2 alone or IL-2 plus TGF-β. Addition of PGE_2_ had few effects on Foxp3 expression in T cells cultured with IL-2 from most donors. However, PGE_2_ suppressed TGF-β-dependent Foxp3 expression in T cells from 3 out of 4 donors (**Figure 5A, B**), suggesting that PGE_2_ may similarly inhibit TGF-β-dependent human iTreg cell differentiation.

**Figure 5.**
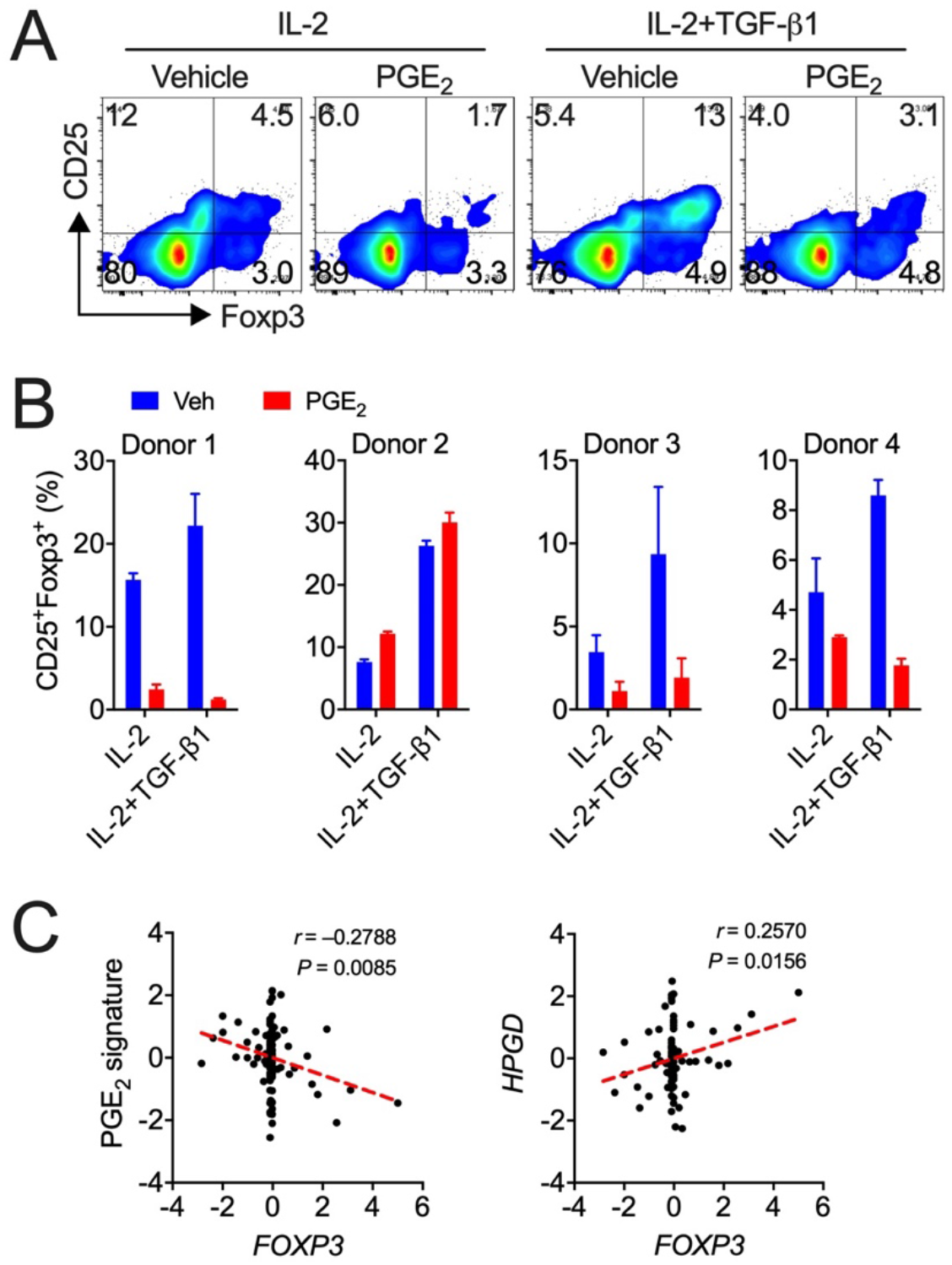
PGE_2_ inhibits human iTreg cell differentiation. (**A**) Representative flow cytometry dot-plot of Foxp3 and CD25 expression in CD4^+^CD45RA^-^ naïve T cells that were isolated from healthy human blood, stimulated with anti-CD3 and anti-CD28, and cultured IL-2 alone or IL-2 + TGF-β1 in the absence or presence of PGE_2_ for 3 days. (**B**) Accumulated percentages of CD25^+^Foxp3^+^ human iTreg cells from four individual donors. (**C**) Microarray gene expression data from human colon biopsies in response to aspirin administration for 2 months in healthy individuals was analysed for the association of the PGE_2_ pathway signature gene expression with that of Treg-related genes. Correlations between the PGE_2_ signature scores or *HPGD* expression levels and Foxp3 gene expression from total tested samples (n=88). Raw gene expression data were retrieved from Gene Expression Omnibus GSE71571. Standardized expression values represent changes of gene expression levels before and after aspirin treatment and then transformed to Z-scores. Each dot represents one sample. Statistical analysis was calculated by two-tailed Pearson correlation coefficients (*r*), and a linear regression-fitting curve is shown as the red dotted line.

We then asked whether the expression levels of PGE_2_ signalling pathway genes were correlated with Foxp3 gene expression in human tissues. We examined a public dataset from a clinical trial which measured gene expression of colon biopsies obtained from healthy individuals before and after administration of aspirin (325 mg/d, daily for 60 days) (Thomas *et al*., 2015). We correlated the changes in mRNA expression of PGE_2_ pathway signature genes (including PGE_2_ synthases: *PTGS1, PTGS2, PTGES, PTGES2, PTGES3*, and receptors: *PTGER2, PTGER4*) before and after aspirin administration with changes in Foxp3 gene expression. Changes in PGE_2_ pathway genes by aspirin treatment were negatively correlated with changes in Foxp3 gene expression (**Figure 5C**). In contrast, changes in expression of *HPGD* (which mediates the metabolic inactivation of PGE_2_ to 15-keto PGE_2_) was positively correlated with changes in *FOXP3* gene expression (**Figure 5C**). These results suggest that the PGE_2_ pathway is associated with down-regulation of Foxp3 gene expression in healthy human gut tissues.

## 4. Discussion

In this report, we show that PGE_2_ directly acts on T cells to abrogate TGF-β signalling and iTreg cell differentiation, especially reducing the CD25^+^Foxp3^+^ subpopulation. Although PGE_2_ did not affect percentages of the Foxp3^+^ population in nTreg cells cultured with TGF-β, it reduced Foxp3 expression at the single cell level. Importantly, PGE_2_ reduced CD25 expression in both iTreg and nTreg cells, indicating that PGE_2_ may lower Treg cell suppressive function. This finding is consistent with other reports, showing that PGE_2_ down-regulates Treg cell responses despite PGE_2_ initially being suggested to facilitate the induction of Foxp3 expression and iTreg cell differentiation *in vitro* (Baratelli *et al*., 2010; Sharma *et al*., 2005; Trinath *et al*., 2013). For example, PGE_2_ inhibited Foxp3 induction and Treg cell proliferation from mouse and human CD4^+^CD25^-^ naïve T cells *in vitro* (B. P. L. Lee *et al*., 2009; H. Li *et al*., 2017; L. Li *et al*., 2019; Sahin & Sahin, 2020). In addition, PGE_2_ was also reported to suppress IL-27-dependent, IL-10-producing type 1 Treg cell differentiation (Hooper *et al*., 2017). Moreover, we have very recently discovered that PGE_2_ suppresses Treg cell accumulation in the intestine through a T cell-independent mechanism (Crittenden *et al*., 2021).

We have found here that PGE_2_ suppression of iTreg cell differentiation is mediated by PGE_2_ receptors EP2 and EP4 *in vitro* and that T cell-specific EP4 deficiency enhanced iTreg cell differentiation *in vivo*, suggesting that the receptor EP2 is dispensable for PGE_2_ suppression of iTreg development *in vivo*. This is consistent with previous findings of the actions of PGE_2_ on Th1 and Th17 cell functions *in vivo* (Yao *et al*., 2009, 2013). Indeed, blockade of EP4 alone sufficiently diminishes Th1/Th17 cell-mediated immune inflammation such as multiple sclerosis, arthritis, and skin inflammation although blockade of EP2 may have additional effects under certain circumstances (Esaki *et al*., 2010; J. Lee *et al*., 2018; Yao *et al*., 2009, 2013). During iTreg cell differentiation, TCR engagement induces T cell activation and production of cytokines such as IL-2 which through activation of STAT5 boosts the induction of Foxp3 expression (Guo *et al*., 2013). Inhibition of STAT5 activity reduced Foxp3 expression during iTreg cell differentiation, which was further repressed by additional PGE_2_, excluding the possibility that PGE_2_ inhibits Foxp3 induction through the TCR-IL-2-STAT5 pathway.

Like actions on other T cell subsets, the inhibition of iTreg cell differentiation by PGE_2_ is also mediated by EP2/EP4-activated cAMP signalling. In Th1 and Th17 cells, cAMP signalling directly induces expression of IL-12Rβ2 and IL-23R, key cytokine receptors for Th1 and Th17 cell differentiation, respectively (J. Lee *et al*., 2018; Yao *et al*., 2013). The mechanism for PGE_2_ inhibition of iTreg cell differentiation is through down-regulation of TGF-β signalling, possibly by reducing expression of TGF-β receptors. Engagement of TGF-β receptors results in phosphorylation of SMAD2/3 and formation of a complex with SMAD4. After translocation into the nuclear, the SMADs complex binds to the CREB/CBP/p300 complex, then in turn regulates transcription responses, including Foxp3 transcription in CD4^+^CD25^-^ T cells (Bodor *et al*., 2007). Indeed, deficiency of CBP and p300 in Foxp3^+^ Treg cells impairs Treg cell stability and suppressive function, resulting in over-activation of effector T cells and autoimmune inflammation (Yujie Liu *et al*., 2014). The transcription factor CREB has also been implicated, as being essential for TCR-induced Foxp3 gene expression (Kim & Leonard, 2007). However, deficiency of CREB in T cells actually decreases Treg cell proliferation and survival and expands Th17 cell response, resulting in exacerbation of T cell-mediated autoimmune inflammation (Wang *et al*., 2017). Thus, PGE_2_-EP2/EP4 signalling activated CREB via cAMP-PKA signalling may contribute to our finding that PGE_2_ prevents TGF-β-dependent induction of Ki-67^+^Foxp3^+^ iTreg cells. Furthermore, the cAMP/PKA/CREB pathway has also been reported to antagonise the TGF-β/SMADs pathway in multiple cell types (Schiller *et al*., 2003). Likewise, we found that PGE_2_ suppressed expression of TGF-β receptor genes (e.g. *Tgfbr1, Tgfbr2*) but upregulated gene expression of TGF-β pathway inhibitors (e.g. *Smad6, Smad7*) in T cells. Lack of TGF-β or its receptors or interruption of TGF-β/SMAD signalling prevents Treg cell development (Yongzhong Liu *et al*., 2008). It is noteworthy that PGE_2_ also inhibited TGF-β/IL-6-induced Th17 cell differentiation although it markedly upregulated IL-23-driven Th17 cell expansion (Yao *et al*., 2009). Additionally, TGF-β-driven Foxp3 strongly down-regulates expression of endogenous inhibitors of TGF-β signalling such as SMAD6 and SMAD7 (Fantini *et al*., 2004), which interference with SMAD-specific gene transactivation. Therefore, down-regulation of TGF-β receptors and upregulation of TGF-β signalling inhibitors by PGE_2_ may collaboratively lead to diminished TGF-β responsiveness during iTreg cell differentiation.

PGE_2_ signalling, especially through the EP4 receptor, is critical for T cell-mediated chronic autoimmune inflammation in numerous organs including skin, joint, brain and intestine etc (Yao & Narumiya, 2019). This was considered to be mediated by promoting inflammatory Th1 and Th17 cells. Our findings in this report suggest that inhibition of Treg cells may be also a mechanism involved in PGE_2_ exacerbation of immune inflammation. Especially, as PGE_2_ suppresses iTreg cell responses *in vivo*, evidenced by transfer of specific EP4 deficiency in T cells reducing Foxp3^+^ iTreg cells in colons of host Rag1^-/-^ mice, which may partially contribute to reduced Th1 cell-medicated colonic inflammation (Yao *et al*., 2013). Furthermore, lack of EP4 in T cells reduced Foxp3^+^ Treg cell accumulation in draining lymph nodes in antigen sensitized mice, which may also contribute, at least in part, to reduced skin inflammation after challenge with the same antigen (Yao *et al*., 2009, 2013).

In addition to previous findings that PGE_2_ indirectly suppresses intestinal Tregs through modulation of the gut microbiota (Crittenden *et al*., 2021), our current results have revealed a direct action of PGE_2_ on T cells to negatively regulate Treg cell differentiation *in vitro* and *in vivo* through EP4 and EP2. These functions of PGE_2_ on Treg cells, together with its positive influences on Th17 and Th1 cells, contribute to facilitation of T cell-mediated tissue inflammation. PGE_2_ inhibition of Foxp3 expression was observed in not only mouse but human T cells during iTreg cell differentiation. Furthermore, negative correlations between the PGE_2_-EP4 pathway and Foxp3 gene expression was observed in healthy human subjects after use of aspirin which inhibits COX activity and PGE_2_ biosynthesis. Thus, therapeutically targeting PGE_2_-EP4 signalling in T cells may be beneficial for treating immune-mediated inflammation, partially due to modulation of Treg cells.

## Acknowledgments

We thank SE Howie and SA Anderton for helpful discussions, warmest support, and providing reagents and the Foxp3^YFP-Cre^ mice; RM Breyer for the EP4-floxed mice; and S Johnston, W Ramsay, M Pattison and F Rossi at the University of Edinburgh QMRI and SCRM flow cytometry facilities for cell sorting and analysis.

